# Dehydration induced *AePer50* regulates midgut infection in *Ae. aegypti*

**DOI:** 10.1101/2023.10.11.561962

**Authors:** Anastasia Accoti, Margaret Becker, Angel Elma I. Abu, Julia Vulcan, Ruimei Yun, Steven Widen, Massamba Sylla, Vsevolod L. Popov, Scott C. Weaver, Laura B. Dickson

**Affiliations:** Department of Microbiology and Immunology, University of Texas Medical Branch, Galveston, TX; Department of Biochemistry and Molecular Biology. University of Texas Medical Branch, Galveston, TX; Faculte des Sciences et Techniques, Universite Cheikh Anta DIOP, Dakar, Senegal; Department of Pathology, University of Texas Medical Branch, Galveston, TX; Center for Vector-borne and Zoonotic Diseases, University of Texas Medical Branch, Galveston, TX; Institute for Human Infections and Immunity, University of Texas Medical Branch, Galveston, TX

## Abstract

In the face of climate change, mosquitoes will experience evolving climates including longer periods of drought. An important physiological response to dry environments is the protection against water loss or dehydration, here defined as desiccation tolerance. Various environmental factors including temperature are known to alter interactions between the mosquito, *Aedes aegypti*, and the arboviruses it transmits, but little is known about how low humidity impacts arboviral infection. Here, we report that a gene upregulated in response to desiccation is important for controlling midgut infection. We have identified two genetically diverse lines of *Ae. aegypti* with marked differences in desiccation tolerance. To understand if the genetic basis underlying desiccation tolerance is the same between the contrasting lines, we compared gene expression profiles between desiccant treated and non-desiccant treated individuals in both the desiccation tolerant and susceptible lines by RNAseq. Gene expression analysis demonstrates that different genes are differentially expressed in response to desiccation stress between desiccation tolerant and susceptible lines. The most highly expressed transcript under desiccation stress in the desiccation susceptible line encodes a peritrophin protein, *Ae*Per50. Peritrophins play a crucial role in peritrophic matrix formation after a bloodmeal. Gene silencing of *Ae*Per50 by RNAi demonstrates that expression of *Ae*Per50 is required for survival of the desiccation susceptible line under desiccation stress, but not for the desiccation tolerant line. Moreover, the knockdown of *Ae*Per50 results in higher infection rates and viral replication rates of ZIKV and higher infection rates of CHIKV. Finally, following a bloodmeal, the desiccation susceptible line develops a thicker peritrophic matrix than the desiccation tolerant line. Together these results provide a functional link between the protection against desiccation and midgut infection which has important implications in predicting how climate change will impact mosquito-borne viruses.

## INTRODUCTION

Arthropod-borne viruses (arboviruses) such as Zika, dengue, and chikungunya viruses transmitted by the mosquito, *Aedes aegypti*, pose a major threat to public health. *Ae. aegypti* has an almost global distribution putting the entirety of the tropics at risk for these viruses. Climate models predict that as global temperatures rise, the risk for arboviral disease will increase causing arboviruses to become a greater global threat, especially in Africa [1, 2]. The ongoing global expansion of Zika [3, 4] and year-round transmission potential by *Ae*. *aegypti* is likely to expand particularly in South Asia and sub-Saharan Africa [1, 5]. Additionally, some models anticipate the climate-driven emergence of dengue and chikungunya at different altitudes and elevations [6–9].

While most predictions about the impact of climate change on vector-borne disease transmission are focused on temperature, little is known about how other climate variables such as humidity will impact vector-borne disease dynamics. Changes in humidity can exist at the micro-or macro level and are generally the result of changes in precipitation. Prolonged periods of reduced rainfall can result in periods of drought or aridity. Aridity can have a significant impact on ecosystems, water resources, and human societies [10]. Like most terrestrial organisms, mosquitoes possess physiological mechanisms to prevent water loss and deal with dehydration mostly through the use of their exoskeleton. The mosquito exoskeleton contains waxes and polysaccharides that prevent water loss across the cuticle. The loss of water can be mitigated through changes in body size, metabolism, composition of cuticular hydrocarbons, and a variety of behavioral and biological adjustments, such as modification in the expression of ion transporter and aquaporin-2 genes [11].

Our understanding of how low humidity impacts the dynamics of vector-borne diseases remains very limited. Water loss has been directly associated with changes in mosquito behavior, leading to an increase in blood feeding rate and potential increases in transmission of West Nile virus (WNV) [12]. Additionally, water loss has been connected to decreased survival and oviposition [13]. Diminished nutritional resources and fecundity have been associated with alterations in both geographic and microgeographic ranges [14, 15].

In this study, we identified two genetically diverse lines of *Ae. aegypti* from Senegal [23] with marked differences in desiccation tolerance and Zika virus susceptibility. We used these lines to probe the genetic basis of desiccation tolerance and identified a gene, *AePer50*, encoding a peritrophin protein that was upregulated in the desiccation susceptible line. Functional validation of *AePer50* confirms its importance in desiccation tolerance in a mosquito genotype dependent manner. We demonstrate that *AePer50* is required to limit midgut infection across arbovirus families in both genotypes. Furthermore, we show that the peritrophic matrix is thicker in the desiccation susceptible/ZIKV resistant line. Together these results provide a functional link between the protection against desiccation and midgut infection which has important implications in predicting how climate change will impact mosquito-borne viruses.

## RESULTS

### Desiccation tolerance is *Ae. aegypti* line-dependent

Survival under acute desiccation was measured in two genetically diverse lines of *Ae. aegypti* originating from Senegal. Establishment and genomic characterization of the lines was done previously [16]. We anticipated differences in survival under desiccation stress between the lines due to the ecology of origin for both lines. The Thiés (THI) line originates from the Northern region of Senegal which exhibits a reduced rainfall compared to the PK10 (PKT) line which originates from the Southern forested region of Senegal. To determine if there are differences in desiccation tolerance between the two lines, we exposed individual females from each line to harsh desiccation (1% relative humidity, RH) and monitored survival for 48 hours. Survival was monitored in individual females not subjected to the harsh desiccation as a control. No difference in survival was detected between the THI line and PKT line under the control condition (Mantel-Cox: p-value = 0.02). Overall, the mosquitoes that were exposed to the harsh desiccation survived less than the control mosquitoes. (Mantel-Cox: p-value < 0.0001). The PKT died at a faster rate compared to the THI line (Mantel-Cox: p-value < 0.0001), with 50% of PKT mosquitoes dying within 24 hours, while 50% of the population survived for 35 hours in the THI line (Figure 1A).

**Figure 1:**
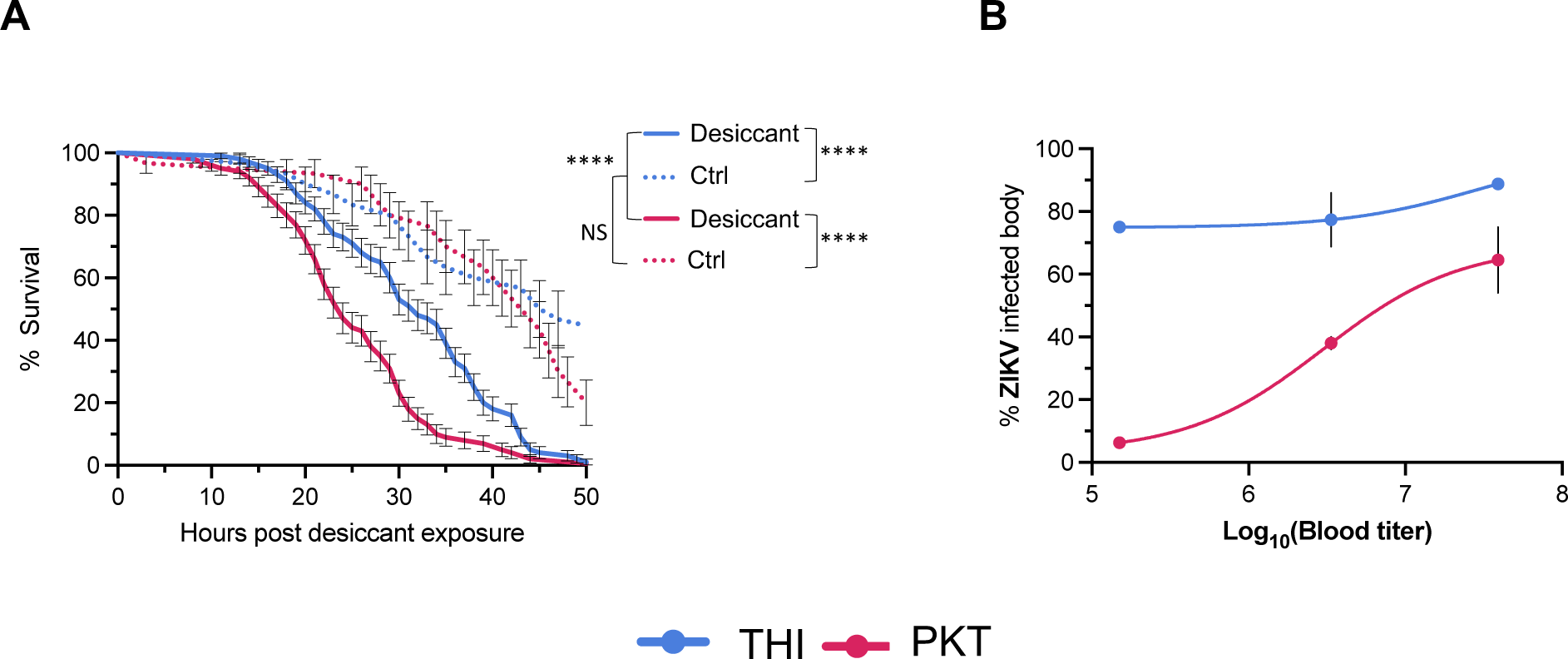
The PKT line is desiccation susceptible and less permissive to Zika virus. **A)** Mosquitoes in single tubes were exposed to acute desiccation stress (RH 1%, 28°C) or ambient conditions (RH 45%, 28°C). Survival was video-recorded for 48 hours and time of death noted. Survival of each line and treatment was compared by a log-rank (Mantel-Cox) test. A total of 100-desiccant-treated-individuals (Desiccant) and 30 untreated from two independent experiments (Ctrl) were used. (THI-Desiccant vs. PKT-Desiccant: p-value: < 0.0001, THI-Ctrl vs. PKT-Ctrl: p-value = 0.02. **B**) Mosquitoes were infected with three different doses of ZIKV (dose I: 7.45×10^7^, dose II: 3.5×10^6^, dose III: 1.5×10^6^) and bodies were collected at 5 days post infectious bloodmeal. Infection rate was determined by reverse transcription-polymerase chain reaction (RT-PCR) with ZIKV specific primers. Doses I and II represent two independent experiments with 10-38 individuals per replicate. Dose III represents a single replicate with 18-32 mosquitoes per treatment. The OID_50_ for THI was 2.246 log10 focus-forming units (FFU)/mL of ZIKV and the OID_50_ for PKT was 7.189 log10 FFU/mL of ZIKV. Overall, differences in ZIKV infection can be explained by ZIKV dose (ANOVA, p-value < 0.0001) and mosquito line (ANOVA, p – value < 0.0001).

Previous studies have demonstrated that lines of *Ae. aegypti* from the North of Senegal are more permissive to ZIKV than lines from the South. To confirm that we observed the same phenotype in the lines, we measured ZIKV infection rates in the THI and PKT lines following exposure to different doses of ZIKV. The proportion of infected individuals was determined 5 days post infection by detecting viral RNA by RT-PCR. In accordance with previous studies, the human-adapted line of *Ae. aegypti* (THI) showed an increased susceptibility to ZIKV compared to the sylvatic line of *Ae. aegypti* (PKT) (Figure 1B). The OID_50_ of the THI line was 2.25 log_10_ focus-forming units (FFU)/ml of ZIKV compared to 7.19 log_10_ FFU/mL of ZIKV in the PKT line. Given that the OID_50_ for THI is outside the range of the dataset, we cannot directly compare the OID_50_ values between lines (THI 95% confidence interval NA-5.21, PKT 6.45-8.38). Overall, differences in ZIKV infection can be explained by ZIKV dose (ANOVA, p-value < 0.0001) and mosquito line (ANOVA, p-value < 0.0001).

### Gene expression profiles under desiccation are *Ae. aegypti* line-dependent

To determine how desiccation stress impacts gene expression in the desiccation tolerant and desiccation susceptible lines, we assessed the global transcriptomes of live whole female bodies 24 hours post 1% desiccation stress. Principal component analysis (PCA) of normalized read counts of 19,804 genes showed that the sequencing libraries clustered by both mosquito line and treatment (Figure 2A) indicating the genetic response to desiccation is different between the two lines. Desiccation stress resulted in both upregulation and downregulation of differentially expressed genes. In the PKT line, a majority of the genes were upregulated in response to desiccation stress, while a similar number of genes were upregulated and downregulated in the THI line (Figure 2B). Strikingly, a single transcript is highly upregulated in the PKT line encoding a peritrophin gene (AAEL002467) previously called *AePer50* [17]. *AePer50* is also upregulated in the THI line, but not to the same magnitude.

**Figure 2:**
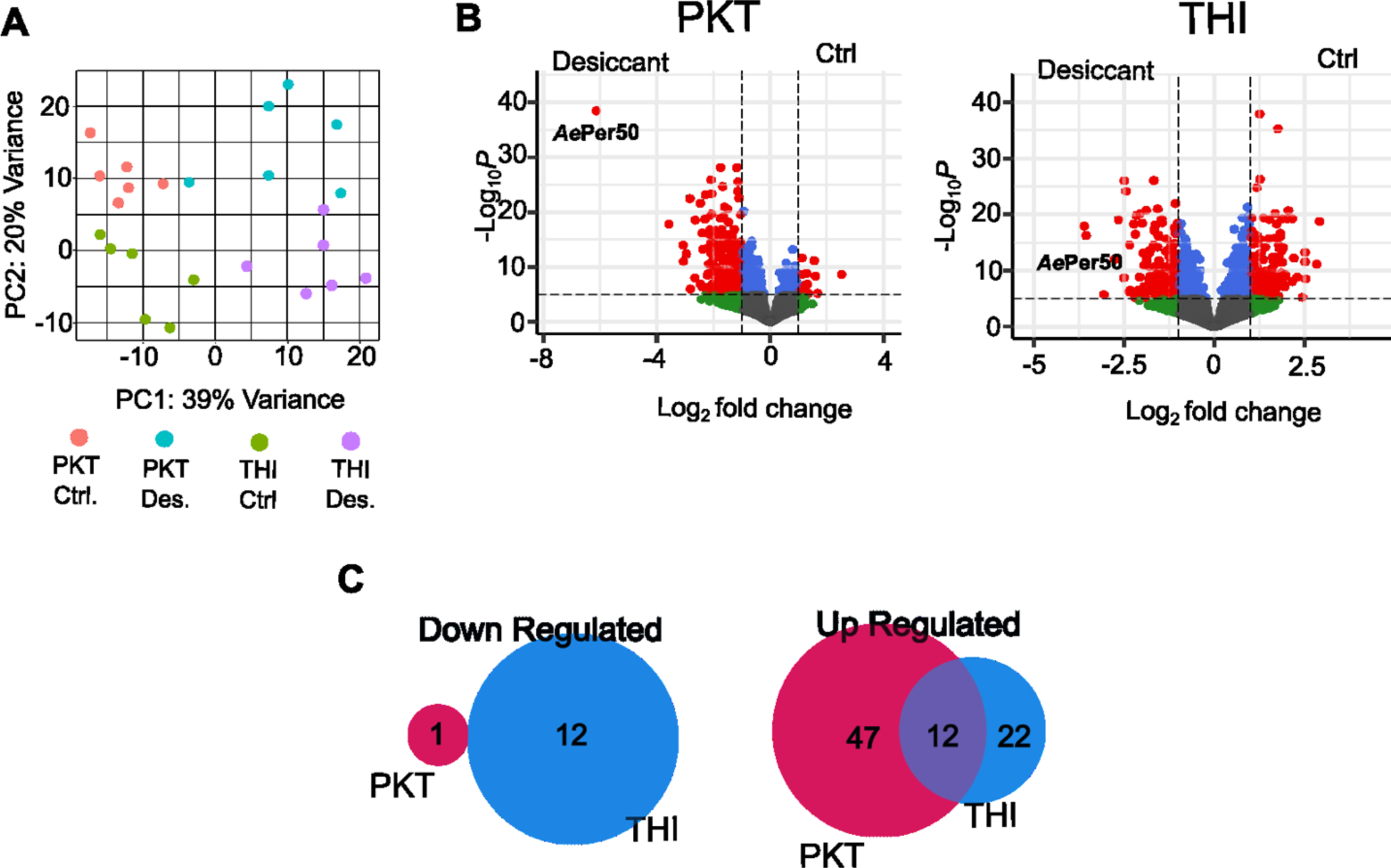
PKT and THI lines show a different genetic response to desiccation stress. Differential expression of genes was determined in response to desiccation stress by RNAseq in the PKT and THI lines. Six libraries were sequenced per treatment containing pools of five individuals. **A**) Principal component analysis of the six sequenced libraries from each treatment. Volcano plot of differentially expressed genes. Gray = not significant, green = log fold change greater than 1, blue = p-value > 0.05, red = log fold change greater than 1 and p-value < 0.05. **C**) Venn diagram showing the overlap of genes either upregulated or downregulated in the THI and PKT lines during desiccation stress with a greater than 2-fold difference.

To better understand the differential transcriptomic response between lines, we performed pair-wise comparative analysis to identify genes differentially expressed between PKT and THI lines. Of the downregulated genes with a log2 fold change greater than two, zero genes were shared between the PKT and THI lines. Twelve genes were upregulated in both the THI and PKT lines, while each line contains upregulated genes specific to the line (Figure 2C and Supplemental Table 1). Gene ontology analysis demonstrates that genes belonging to different biological processes were upregulated in each line (Supplemental Figure 1, Supplemental Table 2).

### *AePer50* expression protects against desiccation in the PKT line

To confirm that *AePer50* expression plays a role in survival under desiccation stress, expression of *AePer50* was knocked down through RNAi mediated gene silencing and survival was measured under desiccation stress in both lines. The THI and PKT lines were injected with dsRNA targeting *AePer50* and dsRNA targeting GFP as control. Single adult females were exposed to 1% RH 72 hours post injection and survival was monitored for 48 hours. Knockdown of *AePer50* did not affect survival under desiccation stress in the THI line (Mantel-Cox: p-value = 0.09) (Figure 3A). In the PKT line, the knockdown of *AePer50* resulted in a reduced lifespan compared to the dsGFP injected control (Mantel-Cox: p-value = 0.0002) (Figure 3B). Knockdown of *AePer50* was confirmed 72 hours post-injection by qPCR (Supplemental Figure 2).

**Figure 3:**
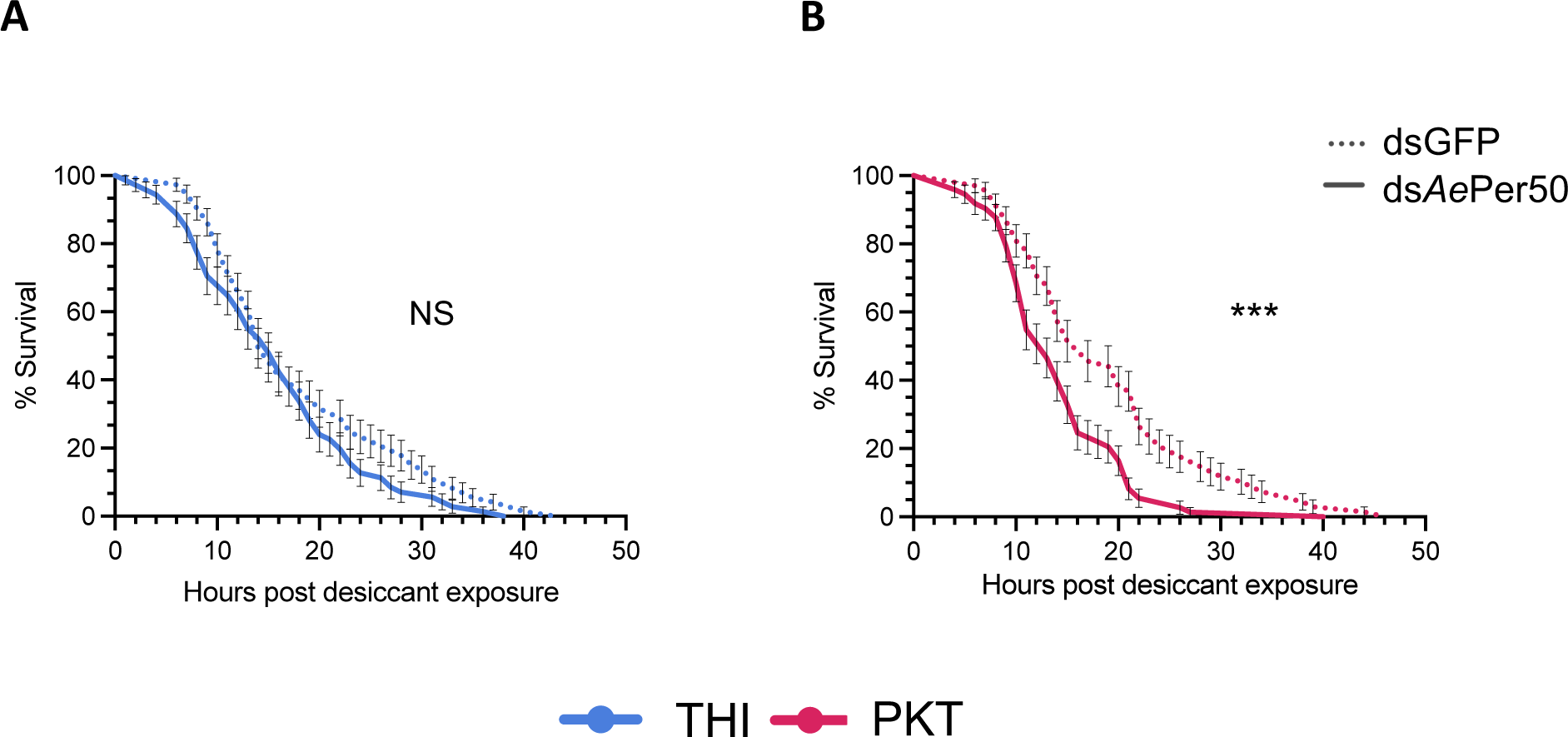
Only PKT relies on *AePer50* for protection against desiccation stress. Survival was measured in individual adult females exposed to acute desiccation stress (RH 1%) 72 hours post injection with dsRNA targeting *AePer50* or GFP as a control for 48 hours. Survival curves were compared by a Log-Rank (Mantel-Cox) test on a total of 90 ds*AePer50* and 50 dsGFP injected mosquitoes. The data represents the mean of three independent experiments, bars represent SEM. (THI ds*AePer50* vs. THI dsGFP: p-value = 0.09, PKT ds*AePer50* vs PKT dsGFP: p-value = 0.0002).

### *AePer50* is expressed in the midgut following a bloodmeal

Expression of the *AePer50* gene has previously been described as being restricted to bloodmeal digestion [17]. To confirm that expression of *AePer50* is also induced following a bloodmeal in our lines, expression of *AePer50* was measured by qPCR following a bloodmeal in each line. In accordance with previously published work, we see induction of *AePer50* expression within four hours following a bloodmeal, and expression diminishing by 12 hours following a bloodmeal in both the THI and PKT lines (Supplemental Figure 3).

### PKT has a wider peritrophic matrix than THI

Peritrophins, and specifically *AePer50*, have been previously characterized for their presence in the peritrophic matrix, a chitinous sac that envelops the blood bolus following a bloodmeal. Given that we see differences in infection rates between the THI and PKT lines, ultrastructural images of blood-fed midguts were taken using transmission electron microscopy to test for any structural differences in the peritrophic matrix between lines. The peritrophic matrix surrounding the blood bolus was measured in 10 different sections of the midguts being sure to avoid areas of breakage area (Figure 4A). The more ZIKV-resistant line (PKT) had a thicker peritrophic matrix (2136.0 ± 964.6 nm) compared to the more susceptible line (THI) (1062.0 ± 547.2 nm) (Mann-Whitney test, p-value of <0.0001) (Figure 5B). To ensure that the increase of thickness was not biased by the overall size of the midgut from each *Ae. aegypti* line, the length and the area of each midgut was analyzed and recorded. Overall, the size of the PKT midguts did not differ from the THI midguts. The PKT midguts had a mean length of 136.5 ± 40.61μm and a mean area of 32405.6 ± 28386.2 μm^2^, compared to the THI which had a mean length of 813.1 ± 260.1 μm and a mean area of 143821.8 ± 129996.9 μm^2^ (Mann-Whitney test, p-value = 0.2).

**Figure 4:**
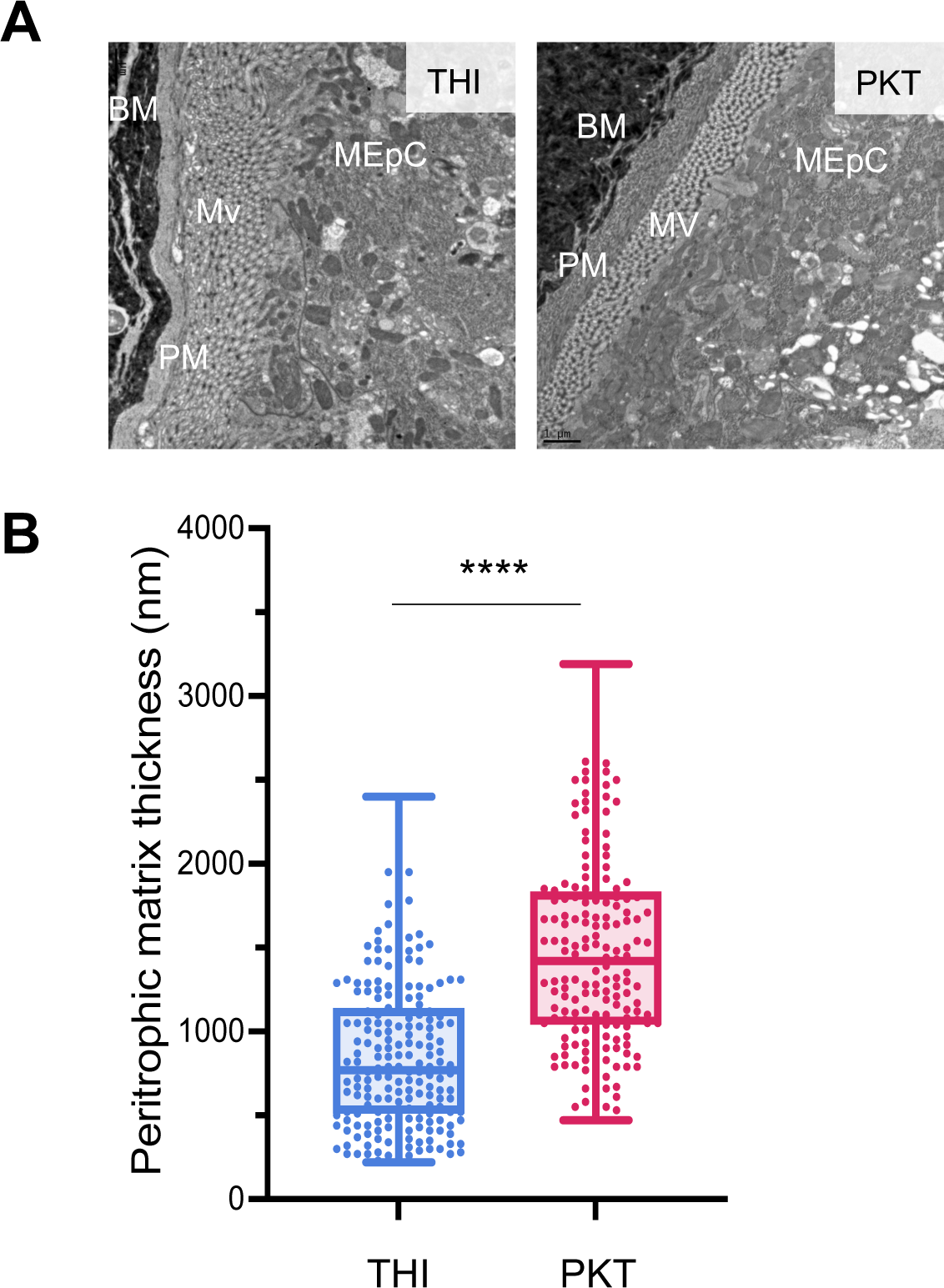
PKT has a ticker peritrophic matrix compared to THI. Electron microscopy was used to capture images of the peritrophic matrix in either the PKT or THI line 22 +/-0.5 hours post bloodmeal**. A)** Representative electron microscopy image of a longitudinal midgut section (80nm) from each line. BM = bloodmeal, PM = peritrophic matrix, Mv = microvillae, MEpC = midgut epithelial cell. **B)** The thickness was measured at a minimum of 10 spots around the entire peritrophic matrix perimeter avoiding the breakage, each dot represent a single measurement. The thickness of the peritrophic matrix was compared between THI and PKT with an unpaired non-parametric t-test Mann Whitney (p-value = < 0.0001).

**Figure 5:**
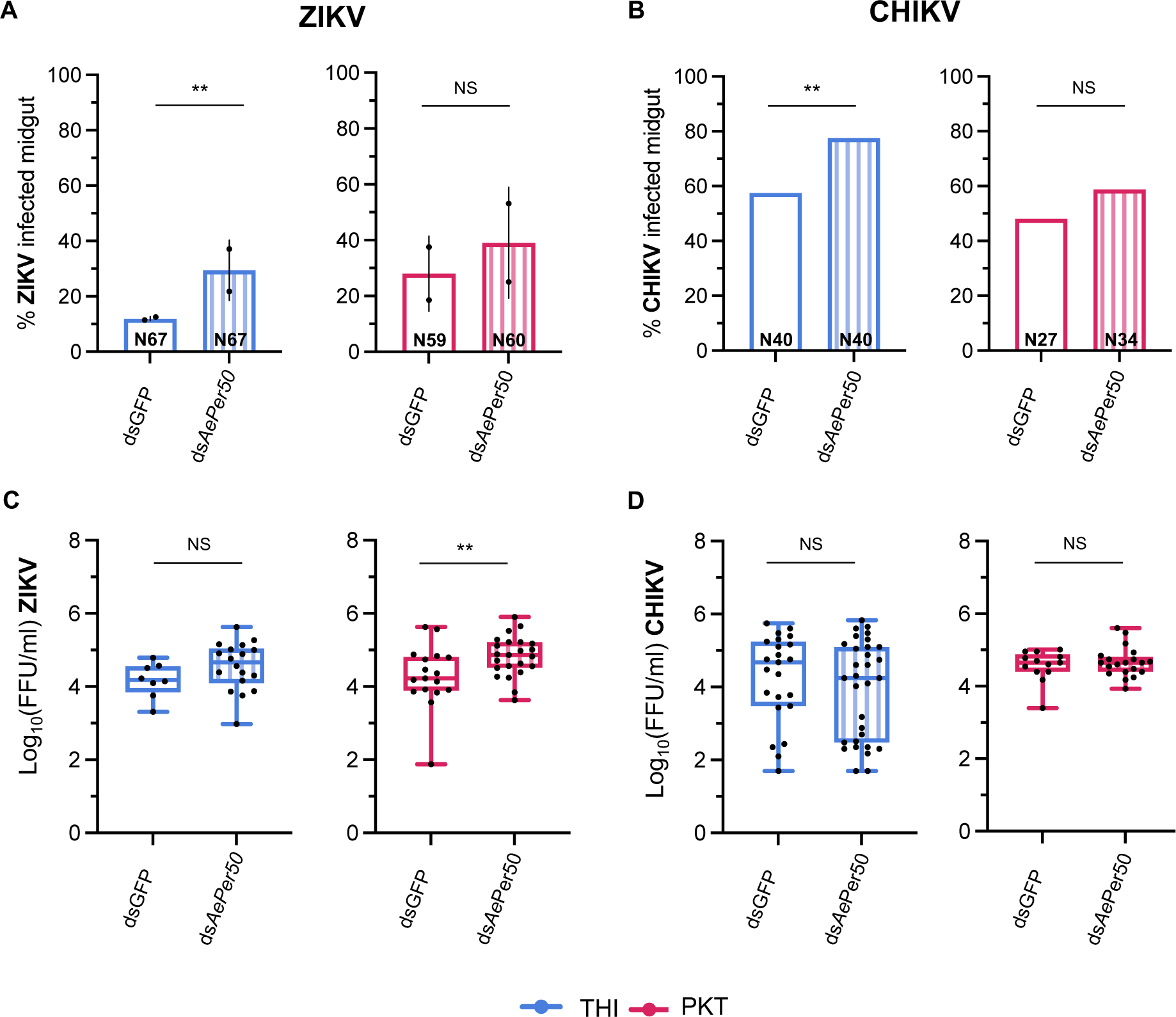
Expression of *AePer50* regulates arbovirus infection. The proportion (**A-B**) of infected *Ae. aegypti* midguts and the amount infectious virus (**C-D**) in *Ae. aegypti* midguts from the THI and PKT lines exposed to ZIKV (**A,C**) or CHIKV (**B,D**) following infection with dsRNA targeting either *AePer50* or GFP as a control. Midguts were dissected 5 days post infectious bloodmeal and proportion of infected midguts and viral titers in the midgut were determined by focus forming unit assay. (**A-B**) Bar graph showing the proportion of infected midguts with either ZIKV or CHIKV following knockdown of *AePer50*. The proportion of infected midguts was compared by Chi-square analysis of a binomial logistic regression as a function of infection, error bars represent the SEM of the proportions. (For ZIKV, THI: dsAePer50 vs. dsGFP p-value = 0.007, PK10: dsAePer50 vs. dsGFP p-value > 0.05). (For CHIKV, THI: dsAePer50 vs. dsGFP p-value = 0.048, PK10: dsAePer50 vs. dsGFP p-value > 0.05). (C-D) Boxplot showing the titers of infectious ZIKV (C) and CHIKV (D) particles in the midgut five days post infection. Each points represents an individual midgut, and the mean is represented by a horizontal line. The error bars represent the 95% confidence interval. Data were analyzed by two-way ANOVA as a function of infection. (For ZIKV, THI: ds*AePer50* vs. dsGFP p-value > 0.05, PK10: ds*AePer50* vs. dsGFP p-value = 0.019). (For CHIKV, THI: dsAePer50 vs. dsGFP p-value >0.05, PK10: ds*AePer50* vs. dsGFP p-value > 0.05). Data represents two independent replicates for the ZIKV infection and one replicate for the CHIKV infection. The total number of individuals tested is displayed on each bar.

### *AePer50* expression controls midgut infection with ZIKV and CHIKV

To determine if *AePer50* plays a role in midgut permissiveness to arboviruses, the THI and PKT lines were injected with dsRNA targeting *AePer50* or GFP as a control. Adult females were offered an artificial bloodmeal containing either ZIKV or CHIKV 72 hours post-injection. Given that *AePer50* is highly expressed following a bloodmeal [17] (Supplemental Figure 3), we measured expression 24 hours following a bloodmeal and confirmed knockdown of *AePer50* (Supplemental Figure 2). Midgut infection rates and viral titers (FFU/ml) in the midgut were assayed five days post virus exposure in dissected midguts. In the THI line, 12% of dsGFP-injected mosquitoes became infected with ZIKV while 30% of ds*AePer50* injected mosquitoes became infected (Chi-Square test, p-value = 0.007) (Figure 4A). Although the ds*AePer50* injected THI mosquitoes had more infectious ZIKV particles in the midgut compared to the dsGFP injected mosquitoes it was not statistically significant (Figure 4B). In the PKT line contrasting phenotypes were observed. Knockdown of *AePer50* resulted in a higher prevalence (37.5 %) of ZIKV infected mosquitoes compared to the control (29.8%) but it was not statistically significant (Figure 4A). Knockdown of *AePer50* resulted in more infectious ZIKV particles in the midgut compared to the control in the PKT line (ANOVA, p-value = 0.019) (Figure 4B).

To determine if the importance of *AePer50* is generalizable across arbovirus families, we challenged mosquitoes with CHIKV following knockdown of *AePer50*. We found a similar trend in the mosquitoes exposed to CHIKV. While a similar trend was observed of a higher proportion of mosquitoes becoming infected following knockdown of *AePer50* compared to the control in both lines (77.5% vs 57.5% in THI, and 58.8% vs 48.1% in PKT) a significant difference in infection rates was only observed in the THI line (THI p-value = 0.048) (Figure 4C). No differences in the number of CHIKV infectious particles were detected in the midgut following *AePer50* knockdown in either line (Figure 4D).

## DISCUSSION

In this study, we use two genetically different *Ae. aegypti* lines which display differences in survival under desiccation stress to probe the genetic basis of desiccation tolerance and identified a dehydration-induced gene that regulates midgut infection with ZIKV and CHIKV. We first identified two lines of *Ae. aegypti* that exhibit differences in survival under desiccation stress. In accordance with previously published results, these lines also display differences in ZIKV infection rates. Total RNA transcriptomics was used to identify the genetic response of both lines to desiccation and we demonstrate that the genetic response differs between the lines. The most highly expressed gene in the desiccation susceptible line encodes a peritrophin protein, *AePer50*. Knockdown of *AePer50* expression confirmed its role in protection against desiccation in the desiccation susceptible line, but not the desiccation tolerant line, further demonstrating the differences between the two lines in responding to desiccation stress.

Because peritrophin proteins are known to be involved in forming the peritrophic matrix, we measured the influence of this gene on midgut infection. Expression of *AePer50* is important for reducing ZIKV and CHIKV replication in both lines indicating the importance of this gene across virus families. Together these data demonstrate that differences in desiccation tolerance is functionally linked to the process surrounding midgut infection which has important implications for predicting how climate change will impact mosquito-borne virus transmission.

An important finding of this study is the differences in the genetic response to desiccation between the two lines. These data suggest that the two lines have evolved different mechanisms to respond to dehydration. This could have arisen from differences in the ecology where these lines originated from. The THI line originates from a drier environment than the PK10 and could have evolved mechanisms to withstand dry environments that the PK10 line has not acquired or lost. Perhaps adapting away from using *AePer50* for desiccation protection is linked to a thinning of the PM.

The involvement of AePer50 during the desiccation stress could be related to its intrinsic ability to stabilize chitin [17], which is part of the insect peritrophic matrix as well as a major component of the insect cuticle [18]. Previous work has shown that *AePer50* is only expressed in the midgut following a blood meal and to a lesser extent after sugar feeding [17]. Additionally, Shao et al. did not detect *AePer50* mRNA in blood-fed or sugar-fed carcasses. Here, RNAseq was done on whole bodies, so we cannot conclude whether *AePer50* expression is occurring in the carcass or the midgut during desiccation stress. AePer50 belongs to the class of peritrophins that have chitin binding motifs containing amino acid sequences consisting of six spaced cysteines referred to as the “peritrophin A domain” [17]. The peritrophin A domain is found in numerous proteins extracted from insect peritrophic matrices [19]. In contrast, chitin-binding motifs in peritrophin proteins are found in the insect cuticle [20]. Given that AePer50 belongs to the class of peritrophins found in the peritrophic matrix, it is interesting that we find it highly expressed in the whole body independent of a bloodmeal and peritrophic matrix formation. One would assume the role of AePer50 protein during desiccation should be related to the midgut and not to the cuticle. The peritrophic matrix is likely to be heavily hydrated as a result of glycosylation of several residues in the linker regions of the peritrophic matrix proteins, and glycosylation is believed to confer protection from proteolytic degradation in addition to conferring resistance to dehydration, infection and physical damage by lubricating the epithelium [17]. Therefore, we could postulate that, specifically in the PKT line, AePer50 has a role in keeping the midgut epithelium hydrated and once this gene is silenced, the PKT line will die sooner than individuals who have full expression of the *AePer50* gene. Further work needs to be done to define the localization and mechanism of AePer50 under desiccation stress.

Reducing the expression of *AePer50* could be increasing midgut infection through disruption of the peritrophic matrix. As we know, one of the main functions of the peritrophic matrix is protecting the mosquito from pathogen invasion [21–23]. Secretion of the peritrophic matrix in *Ae. aegypti* begins around 4 hours after the bloodmeal and it will become separated from the microvilli at 8 to 12 h after blood feeding [24]. Our data indicate the level of *AePer50* expression is important for both midgut infection and replication of ZIKV as well as midgut infection of CHIKV. The importance of this gene in controlling viral infection across viral families suggests that *AePer50* plays a structural role in the peritrophic matrix that is important for controlling access to the midgut epithelium, as opposed to a mechanism that involves viral tropism.

Here we identified the differential thickness of the peritrophic matrix between the THI and PKT lines. Specifically, we found that the desiccation susceptible/ZIKV refractory line has a thicker peritrophic matrix than the desiccation tolerant/ZIKV permissive line. It is tempting to speculate that the differences in ZIKV infection rates between the THI and PKT lines are due to the width of the peritrophic matrix. This is supported by the fact that we can see a significant reduction in ZIKV infection rates in the THI upon silencing *AePer50*, but not in the PKT line. Perhaps this is due to the fact that the peritrophic matrix of THI is thinner to start and we are able to reduce the thickness to a meaningful level with our gene silencing assay, which is not the case with the thicker PKT line. Given that we see a trend in increased infection rates after silencing *AePer50* in the PKT line, it is likely that a higher degree of gene silencing would allow for complete thinning of the peritrophic matrix and a significant increase in midgut infection rates.

We also observed the role of *AePer50* in rates of viral replication in the midgut. This time only a significant difference between dsGFP and ds*AePer50* treated mosquitoes was observed in the PKT line. This suggests that a function of AePer50 outside of the thickness of the PM regulates rates of viral replication. These data indicate that the lines have different mechanisms to control rates of viral replication and that the overall ability of the midgut to become infected at all is a distinct phenotype from rates of viral replication. Here we observe that the ability to block initial infection of the midgut epithelium does not translate to a reduction in rates of viral replication. Once the virus has invaded the midgut epithelium, it seems that other factors control how much the virus can replicate.

AePer50 could be impacting midgut infection through other mechanisms besides establishing the thickness of the peritrophic matrix. Several factors have been shown to influence the ability of a virus to escape from the blood bolus to the epithelial cells, referred to as the midgut infection barrier [24]. The viral particle likely needs to enter the epithelial cells before the formation of the peritrophic matrix. Although we observed a thicker peritrophic matrix in the PKT line, there are other possible explanations for why this line is more refractory than the THI line. Differences in timing of *AePer50* expression and peritrophic matrix formation, the composition of the peritrophic matrix (how much *AePer50* protein is in the peritrophic matrix), or the permeability/stability of the peritrophic matrix could also impact the ability of the virus to reach the midgut epithelium. The thicker peritrophic matrix in the PKT line could be a result of differential timing of formation, resulting in an earlier and longer PM secretion that results in a wider peritrophic matrix. We did not investigate the PM composition in the *Ae. aegypti* lines we used in this study, and this is worthy of further investigation. Perhaps the PKT line has a higher density of the *AePer50* protein in the peritrophic matrix resulting in a stronger barrier for arboviral infection.

Overall, this study demonstrates a functional link between dehydration tolerance and midgut infection which has important implications for predicting how climate change will impact mosquito-borne virus transmission.

## MATERIALS AND METHODS

### Data accessibility

Data, protocols, code, materials will be accessible through NCBI and Git Hub.

### Aedes aegypti mosquitoes

Colonies of *Ae. aegypti* used in this study originated from a natural population of Thiés (THI) and PK10 (PKT) originally sampled respectively in the area of Thiès and the PK10 forest in Senegal [16]. Colonies were made by collecting eggs from each colony using ovitraps as described by Rose et al. [16]. The mosquitoes of the study belong to generation 9 or 10 depending on the experiment. Mosquitoes were reared under standard insectary conditions consisting of 28°C and 70% +/-10% relative humidity with a 12h:12h light-dark cycle. Eggs were hatched in deionized water and larvae were fed fish food. Adults were held in Bugdorm cages with constant access to 10% sucrose until being used for specific experimental procedures.

### Desiccation assay

To determine the acute desiccation tolerance, the method previously described by Ayala et al. 2019 [25] was adapted as follows. Individual 4-5 days old female mosquitoes from both the THI and PKT colonies were cold-anesthetized and placed into plastic *Drosophila* vials (Flystuff Narrow Drosophila Vials, Polystyrene) and plugged with a foam stopper (Flystuff 59-200 Droso-Plugs^®^) approximately 2 cm below the rim of the vial. After all mosquitoes were added to the vials, a desiccation agent (Drierite™ Mesh size 8) was added on top of the foam stopper and parafilm was used to seal the vial. The amount of desiccant agent added was normalized by using a scoop with a volume of approximately 6 g of desiccant agent. Controls had no desiccant agent added and were sealed with parafilm as the treated mosquitoes. Neither the controls nor the desiccant treated mosquitoes had access to sugar or water. The desiccant agent reduced the relative humidity within the vials to <1% and the control vials resulted in a RH of 45% (Onset HOBO External Temp/RH Data Logger). The experiment started upon sealing the vials and was terminated after 48 H. Vials were placed on shelves with approximately 50 vials per shelf inside an incubator (Thermo Fisher-Precision) set to 28°C with a 12h light-dark cycle. To monitor the status of the mosquitoes, home security cameras (Wyze Cam v3) were used. Each camera captured the activity of approximately 50 mosquitos with an HD resolution of 1920 x 1080p at 20Hz during the light cycle and 15Hz during the dark cycle. The cameras use infrared light to record during the dark cycle. The time of death for each individual mosquito was estimated by watching footage of the beginning of each hour over the study period. Mosquitos were considered dead when they were knocked down or otherwise immobile. For every line, a total of 50 individuals for the desiccant and 15 for the control were used in two separate experiments. The statistical significance of survival curves was set to the conventional α < 0.05 level, calculated with a log-rank Mantel-Cox analysis and using GraphPad Prism software, version 10.

### Vector competence

THI and PKT females 4-5 days old were starved for 24 hours before the infectious blood meal. Vector competence assays were performed in an Arthropod Containment level 2 facility (ACL-2). Briefly, mosquitoes were experimentally exposed to different titers of wild-type Zika virus Cambodia isolate (FSS 13052) received from the World Reference Center for Emerging Viruses and Arboviruses at UTMB. The virus stock was diluted in cell culture media (Dulbecco’s modified Eagle Medium (DMEM) with the addition of 1.5% heat-inactivated FBS, 1% penicillin/streptomycin) and 30 μl of 7.5% sodium bicarbonate to reach a dose of 7.5×10^7^ (dose I), 3.5 x 10^6^ (dose II)and 1.5x 10^5^ (dose III) FFU/ml. One volume of virus suspension was mixed with two volumes of defibrinated sheep blood (Colorado Veterinary Product) washed three times in 1X PBS and 60 μl of 100 mM adenosine 5′-triphosphate. After gentle mixing, 2 ml of the infectious blood meal was added to Hemotek membrane feeders (Hemotek Ltd.) covered with a piece of desalted porcine intestine as a membrane and maintained at 37°C. After feeding, fully engorged females were sorted into 1-pint cardboard cups and maintained under controlled conditions (28 ± 1°C; relative humidity, 75 ± 5%; 12h:12h light/dark cycle) in a climatic chamber for 5 days. After 5 days of incubation, whole bodies of ZIKV-exposed mosquitoes were harvested in 2 ml screw-lid vials and homogenized in 300 µl of crude RNA extraction buffer/Squash buffer (10 mM Tris HCl, 50 mM NaCl, 1.25 mM EDTA, fresh 0.35 g/L proteinase K) in a Precellysis homogenizer for 3 rounds of 20 seconds with a 30 second of pause every round, after homogenization, 200 µl of each sample was transferred to a 96-well plate and incubated at 56 °C for 5 minutes followed by 98 °C for 10 minutes. cDNA was produced from 5 µl of each sample using M-MLV reverse transcriptase (Invitrogen) and random hexamers by the following program: 10 min at 25°C, 50 min at 37°C and 15 min at 70°C. The cDNA (2.5µL) was amplified by PCR carried out with DreamTaq DNA polymerase (Thermo Fisher) and specific ZIKV primers (forward: 5’-GTATGGAATGGAGATAAGGCCCA-3’, and reverse: 5’-ACCAGCACTGCCATTGATGTGC-3’) [26]. Amplicons were visualized by gel electrophoresis on a 2% agarose gel. A total of 64 and 58 females were used for THI and PKT respectively. The proportion of ZIKV-infected females was analyzed by Chi-square analysis on a binomial logistic regression as a function of treatment and colony in R. For dose I and dose II, two independent biological replicates were conducted, with a total of 70 individuals for THI and 43 for PKT and 53 individuals for THI and 24 for PKT, respectively. For dose III, a single replicate was carried out, with 32 individuals for THI and 18 for PKT.

### RNA sequencing

THI and PKT females 4-5 days old were exposed for 24 hours to 1% of relative humidity (RH), following the treatment, whole bodies were homogenized in 800mL of TRIzol (Invitrogen) and RNA was extracted with the phenol-chloroform method following the manufacturer’s instruction. The final resuspension of RNA was in 20µL of nuclease-free water.

The UTMB Next Generation Sequence (NGS) core laboratory assessed RNA concentrations and quality using a Nanodrop ND-1000 spectrophotometer (Thermofisher, Waltham) and an Agilent Bioanalyzer 2100 (Agilent Technologies, Santa Clara, CA). PolyA+ RNA was purified from ∼100 ng of total RNA and sequencing libraries were prepared with the NEBNext Ultra II RNA library kit (New England Biolabs) following the manufacturer’s protocol. Libraries were pooled and sequenced on an Illumina NextSeq 550 High Output flow-cell with a paired-end 75 base protocol.

STAR alignment software, version 2.7.10a, was used to build a genome index and map the reads. The index was built from the *Ae. aegypti* LVP_AGWG genome and annotation files downloaded from VectorBase.org, release 59. Reads were mapped with the parameters recommended for the ENCODE consortium and reads mapping to genes were quantified with the STAR–quantMode GeneCounts option. Differential gene expression was estimated with the R v4.1.3 DESeq2 software package, version 1.32.0, following the vignette provided with the package. Log fold changes were moderated with the lfcShrink function using the ashr package.

#### Gene enrichment analysis

Significant genes with at least a 2-fold difference in expression were analyzed for functional enrichment by submitting the gene lists to VectorBase Release 65. Enriched gene ontology (GO) terms were identified using the *Ae. aegypti* LVP_AGWG background and clustered with REVIGO at a medium similarity using Drospila melanogaster as the closest species in the database.

### *AePer50* gene expression time course

To monitor the line-specific *AePer50* gene expression, 4-5 day old female *Aedes aegypti* THI and PKT lines were blood fed a non-infectious bloodmeal and midguts were collected at different time points. Mosquitoes were starved 24 hours before the blood feeding. Feeders were prepared as described in a previous section on the day of the feed. Mosquitoes were fed for 30 minutes and the fully engorged were quickly sorted on ice. Midguts were dissected at 4, 8, 12, 24, and 48 hours post bloodmeal, and midguts from unfed females were used as a control.

Midguts were placed in vials containing 0.1 mm glass beads and 200 μL of TRIzol (Invitrogen), homogenized after collection and stored at −80°C until processing. RNA was extracted using a Quick-RNA Miniprep kit (Zymo). *AePer50* reverse transcription (RT) and qPCR reactions were performed using GoTaq® 1-Step RT-qPCR System (Promega), with 10 µl of 2X qPCR Master Mix, 1 µl of 10 µM primer and 3 µl of cDNA and 0.4 µl of RT enzyme, made up to a total reaction volume of 20 µl with nuclease-free water. Primers used were either *AePer50* primers (FWD-TCATCCTCACCTTCGCCTAC, REV-AAGCTCTGTCGTCGTTGTGGG) or S7 primers (FWD-GCA GAC CAC CAT TGA ACA CA, REV-CAC GTC CGG TCA GCT TCT TG) as a housekeeping gene. Positive control and no template control were added to every qPCR run to assess for contamination and cross primer-dimer formation. The *AePer50* qPCR was performed on a QuantStudio 6 Real-Time PCR System (Applied Biosystems), with cycling conditions as follows: initial denaturation at 95 °C for 20 s and 45 cycles at 95 °C for 10 s and 56 °C for 30 s, and the melting curve 95°C for 10 min, 65°C to 95°C with an increment of 0.5°C every 0.05 seconds. All qPCR reactions were performed in duplicate, and the expression levels of target genes were normalized to the levels of ribosomal protein S7. The fold change in the gene expression was calculated according to the standard ddCT method, using the unfed midgut as the control. For every mosquito line, a total of 12 midguts per time point were processed.

### Gene silencing assay

For producing *AePer50* line-specific dsRNA, total RNA was extracted with Quick-RNA Miniprep kit (Zymo) from 10 whole nonblood fed (NBF) *Ae. aegypti* females according to manufacturer’s instructions including DNase treatment (Zymo). cDNA was generated from 1 μg of total RNA using M-MLV reverse-transcriptase (Invitrogen) and random hexamers (Invitrogen). cDNA was used for the production of the dsRNAs targeting *AePer50* THI-and PKT-gene and the plasmid template *pUC57* (Addgene) was used for dsGFP (used as control dsRNA). dsRNAs were produced as previously described [27] using primers with T7 RNA polymerase promoter sequence [28]. AePer50 dsRNA primers were designed using the E-RNAi web service (https://www.dkfz.de/signaling/e-rnai3/0).

RNA interference assays (RNAi-based gene silencing) were conducted as previously reported [27]. Briefly, 69 nl of dsRNA (3 µg/µl) re-suspended in water was injected into the thorax of cold-anesthetized 3-4 day old female mosquitoes using a Nanoject III (Drummond Scientific).

To evaluate the knockdown efficiency, we measured *AePer50* expression at two time points, whole bodies at 72 H post injection and midguts at 96 H post injection/24 H post bloodmeal. We checked whole bodies to confirm knockdown in our desiccation survival assay, and checked midguts 96 H post injection/24 H post bloodmeal to ensure *AePer50* was still being silenced despite induction of the gene following a bloodmeal. RNA extraction, RT, and qPCR were performed as described in the previous section and the average of fold changes were used to calculate the percentage of knockdown. Knockdown efficiency was determined by normalizing the fold change of *AePer50* relative to S7 housekeeping genes in the ds*AePer50* injected mosquitoes to dsGFP injected mosquitoes.

### Arboviral infection and focus forming assay on *AePer50* injected *Ae. aegypti*

Mosquitoes 72 H post dsRNA injection, performed as described in the previous section, were infected with the wild-type Zika virus Cambodia isolate (FSS 13025) or the wild-type Chikungunya virus isolate (FSS 37997, Senegal, mosquito, 1983, West African lineage) received from the World Reference Center for Emerging Viruses and Arboviruses at UTMB. The infectious bloodmeal was prepared as described before, for the Cambodia infection, two replicates were performed yielding a titer of 2.3 X 10^6^ and 3.5 X 10^6^ FFU/ml for the THI line and of 4 X 10^6^ and 3.6 X 10^6^ FFU/ml for the PKT line. For the CHIKV infection the blood meal titer was 7 x 10^6^ FFU/ml for both *Ae. aegypti* lines. To assess the infection rate of the midguts, a focus-forming assay in Vero cells was performed. Briefly, midguts were homogenized individually in vials containing 0.1 mm of glass beads and 200 µL of Vero cell media, which consists of DMEM with 1% penicillin/streptomycin (Pen-Strep), 1% antibiotic and antimycotic (GIBCO), 5% heat-inactivated fetal bovine serum (FBS). Samples were stored at −80°C until being processed. Vero cells were seeded in 24-well plates and incubated for 48 hours to reach confluency. Each well was inoculated with 200µl of midgut homogenate in 10-fold dilutions (from 10^1^ to 10^4^) and incubated at 37°C (5% CO^2^) for 1 hour, rocking every 15 minutes. Infected cells were overlaid with OPTI-MEM media supplemented with 1.25% carboxymethyl cellulose, 5% FBS, and 1% Pen-Strep. After three days of incubation at 37°C, infected cells were fixed with 10% formalin for at least 1 hour and were washed three times in 1X PBS. Approximately 500µl of blocking solution (5% w/v non-fat powdered milk in 1X PBS) was added to each well and plates were rocked for 30 minutes. The blocking solution was discarded and 200µl of primary antibody solution (obtained from the World Reference Center for Emerging Viruses and Arboviruses-WRCEVA at UTMB) diluted 1:1000 in blocking solution was added to each well and plates were placed on plate rocker overnight. The primary antibody solution was discarded, and plates were washed three times with 1X PBS prior to the addition of 200µl of secondary antibody (peroxidase-labeled goat anti-mouse IgG human serum KPL-474-1806) solution diluted 1:2000 in blocking solution. Plates were placed on plate rocker for 1 hour. The secondary antibody solution was discarded, and plates were washed three times with 1X PBS. To develop visible foci, 100µl of TrueBlue peroxidase substrate (KPL 5510-0050) was added to each well, and plates were placed on the plate rocker until foci could be seen, around 10 min. Plates were washed with deionized water and FFU was counted with the help of a light. Focus-forming units were Log_10_ transformed to represent the concentration of infectious ZIKV particles detected in *Ae. aegypti* midguts. For the ZIKV experiment, two independent experiments were performed and per each knock down treatments (dsGFP and dsAePer50) THI had 32 ± 3 for and PKT had 32 ± 4. For the Chikungunya experiment, one experiment was performed and per each treatment THI had 40 and PKT had 34 ± 7. Data were analyzed by two-way ANOVA as a function of infection in R.

### Peritrophic Matrix Electron Microscopy

For ultrastructural analysis of ultrathin sections THI and PKT females of 4-5 days old were starved for 24 hours before the blood meal, that consists in 2 ml of de-fibrinated sheep blood (Colorado Veterinary Product) washed three time in 1X PBS with the addition of 60 μl of 100 mM adenosine 5′-triphosphate. Only fully engorged mosquitoes were sorted into 1-pint cardboard cups and maintained under controlled conditions (28 ± 1°C; relative humidity, 75 ± 5%; 12:12-hour light/dark cycle) in a climatic chamber for 22 ± 0.5 hours when they were carefully dissected in 1X PBS, avoiding pulling the midgut. After collection, each midgut was fixed in 1 ml of a mixture of 2.5% formaldehyde prepared from paraformaldehyde, 0.1% glutaraldehyde, 0.01% picric acid and 0.03% CaCl_2_ in 0.05 M cacodylate buffer pH 7.3. They were washed in 0.1 M cacodylate buffer, post-fixed in 1% OsO_4_ in 0.1M cacodylate buffer pH 7.3 for 1 h, washed with distilled water and *en bloc* stained with 2% aqueous uranyl acetate for 20 min at 60 °C. Then they were dehydrated in ascending concentrations of ethanol, processed through propylene oxide and embedded in Poly/Bed 812 (Polysciences, Warrington, PA). Before cutting ultrathin sections, 1 μm semi-thin sections were cut and stained with toluidine blue. Semi-thin and ultrathin sections were cut on Leica EM UC7 ultramicrotome (Leica Microsystems, Buffalo Grove, IL), stained with lead citrate and examined in a JEM-1400 (JEOL USA, Peabody, MA) transmission electron microscope at 80 kV. Digital images were acquired with a bottom-mounted CCD camera Orius-SC200-1 (Gatan, Pleasanton, CA). The midgut was cut along the longitudinal plane and pictures were taken around the whole midgut perimeter in areas without peritrophic matrix detachment. A minimum of 10 different sections along all midguts were chosen to take multiple measurements with Gatan Microscopy Suite software (version 1.85.1535). To avoid bias related to the size of the midgut, the peritrophic matrix size measurements were taken together with their respective midgut length and area, measured from a 0.5 μm midgut section, stained with 1% toluidine blue and measured with NIS-Element BR Viewer software (version 4.4). For every mosquito line, three independent experiments were performed, one midgut per replicate. The mean width of the peritrophic matrix was compared between the THI and PKT lines with a Mann Whitney unpaired T-test in Graphpad Prism (Version 10).

## ACKNOWLEDGEMENTS

We would like to thank Jiehua Zhou and Ruimei Yun of the UTMB Insectary Core. We would like to thank Dr. Noah Rose and Dr. Caroline McBride for sharing the mosquito colonies with us. LBD was supported by UTMB start-up funds and U01AI151801 West African Center for Emerging Infectious Diseases. MB was a recipient of the T32 predoctoral fellowship. NIAID T32 Emerging and Tropical Infectious Diseases Training Program (AI007526 to Lynn Soong).

---

**Supplemental Figure 1:**
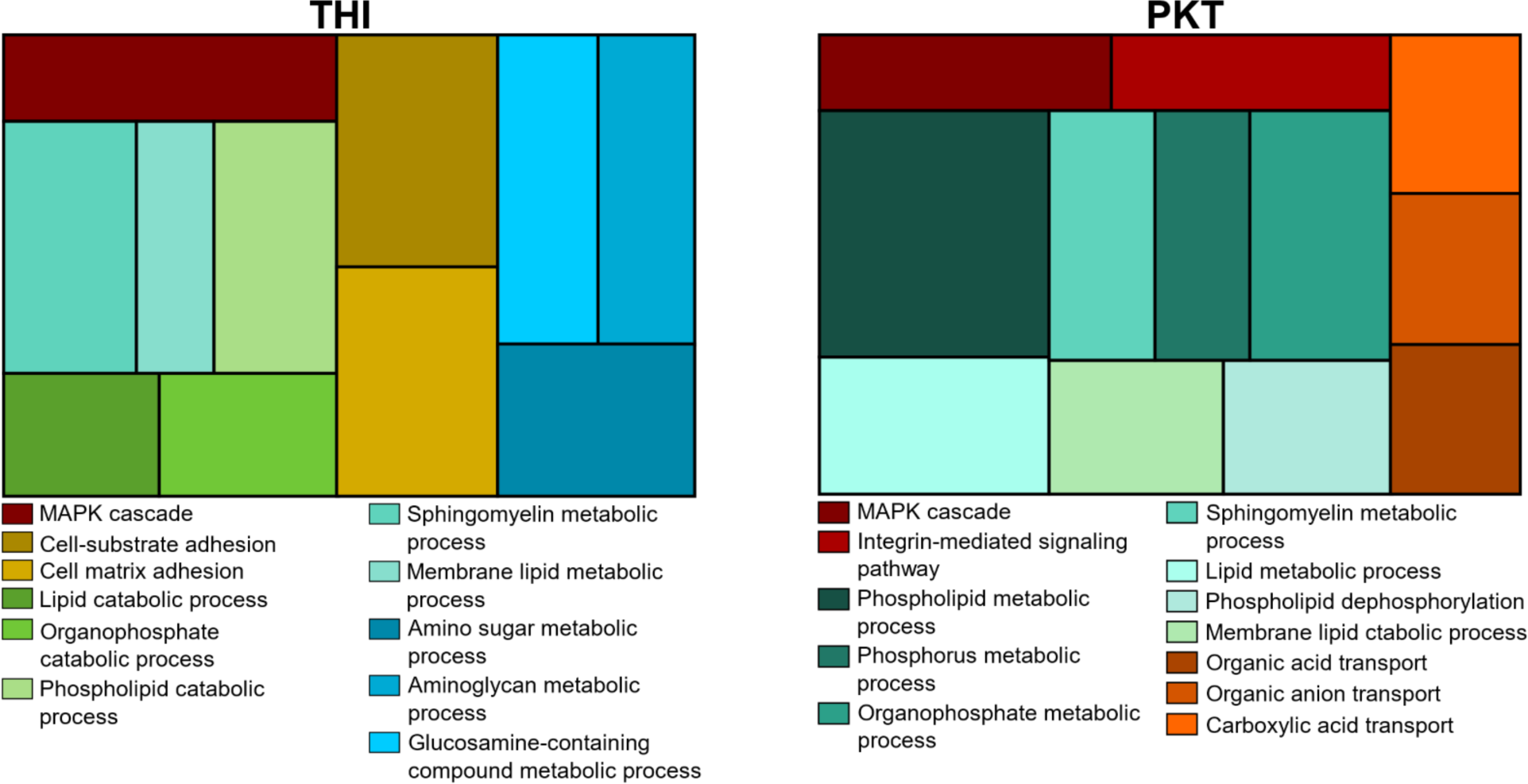
Gene ontology. TreeMap visualization of biological processes enrichment for upregulated genes with a log fold change greater than 2. Color grouping is based on higher level GO term annotation. The size of the rectangle is proportional to the number of genes in the list out of the total number of genes with similar roles in the *Ae. aegypti* genome.

**Supplemental Figure 2:**
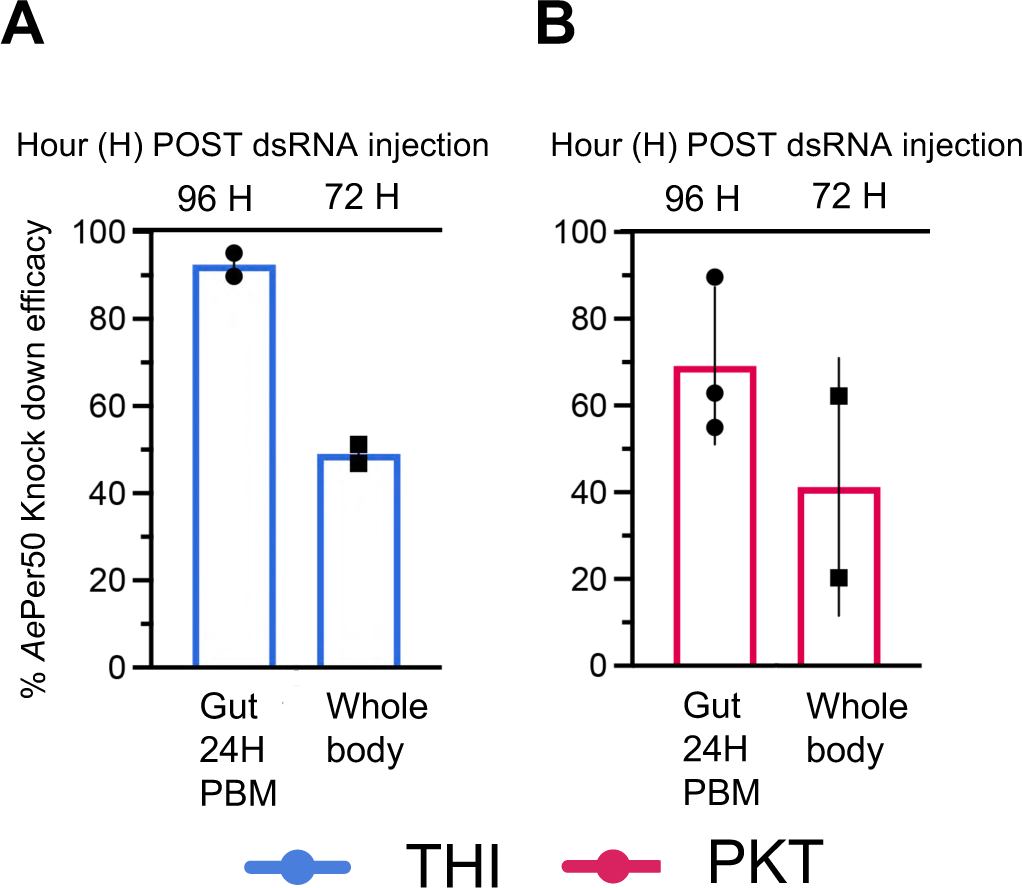
Knockdown efficiency. Mosquitoes were injected with 69 nl of dsGFP or dsAePer50 (3 µg/µl) and knockdown was confirmed in the timepoints and tissues used in our assays. Total RNA was extracted from whole bodies 72 hours post injection as in the desiccation survival assay, and 96H post injection/24H post bloodmeal as in the viral challenge assay. AePer50 transcripts and hourskeeping gene S7 transcripts were quantified by Rt-qPCR. Knockdown efficiency was determined by normalizing the fold change of AePer50 relative to S7 housekeeping genes in the ds*AePer50* injected mosquitoes to dsGFP injected mosquitoes.

**Supplemental Figure 3:**
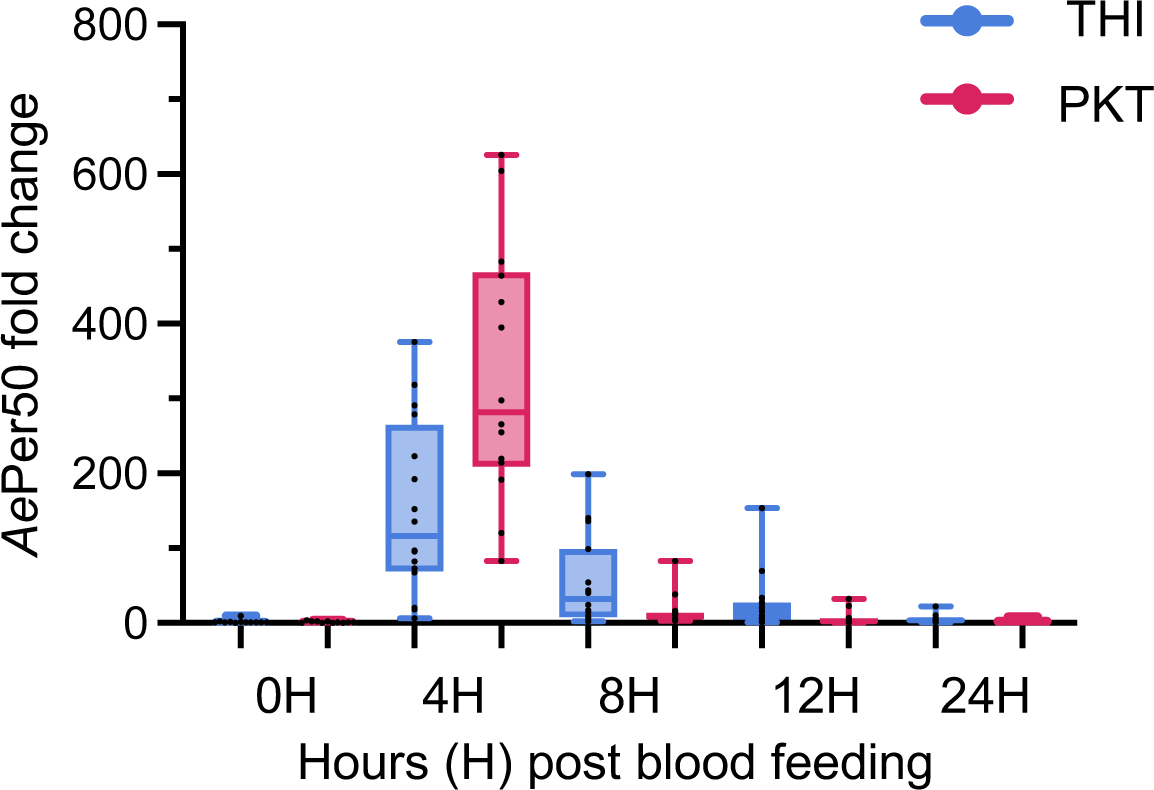
*AePer50* expression time course after a blood meal. Mosquitoes were offered a blood meal and midguts were dissected at 4, 8, 12 and 24 hour post blood feeding, as a control unfed midgut were used. The fold change in the gene expression was calculated according to the standard ddCT method, using the unfed midgut as the control. For every mosquito line a total of 12 midguts per time points were processed.

